# Cytoprotection by a naturally occurring variant of ATP5G1 in Arctic ground squirrels

**DOI:** 10.1101/2020.01.31.929018

**Authors:** Neel S. Singhal, Meirong Bai, Evan M. Lee, Shuo Luo, Kayleigh R. Cook, Dengke K. Ma

**Author notes:** These authors contributed equally (Neel S. Singhal, Meirong Bai). **Corresponding Author:** Dengke K. Ma.

## Abstract

Many organisms, from anaerobic bacteria to hibernating ground squirrels, have evolved mechanisms to tolerate severe hypoxia or ischemia. In particular, the arctic ground squirrel (AGS) has been shown to be highly resilient to ischemic and reperfusion injuries, demonstrating an ability to withstand metabolic stress under hibernation conditions. Although physiological adaptations are critical to ischemic tolerance in AGS, little is known about cellular mechanisms underlying intrinsic AGS cell tolerance to metabolic stressors. Through cell survival-based cDNA expression screens and comparative genomics, we have discovered that in AGS, a cytoprotective variant of ATP5G1 helps confer improved mitochondrial metabolism and cell resilience to metabolic stress. *ATP5G1* encodes a proton-transporting subunit of the mitochondrial ATP synthase complex. Ectopic expression in mouse cells and CRISPR/Cas9 base editing of the endogenous AGS locus revealed causal roles of one AGS-specific amino acid substitution (leucine-32) in mediating the cytoprotective effects of AGS ATP5G1. We provide evidence that AGS ATP5G1 promotes cell resilience to stress by modulating mitochondrial morphological change and metabolic functions. Thus, our results identify a naturally occurring variant of ATP5G1 from a mammalian hibernator that causally contributes to intrinsic cytoprotection against metabolic stresses.

## Introduction

Arctic ground squirrels (AGS) survive harsh winter environmental conditions through hibernation. By virtue of their profound ability to suppress metabolism and core temperature, with body temperatures dropping below 0°C, AGS are known as ‘extreme’ hibernators (1). Hibernation in AGS can last 7 months and is characterized by drastic (>90%) reductions in basal metabolic rate, heart rate, and cerebral blood flow (2). Curiously, hibernation is interrupted periodically by interbout arousal (IBA) episodes in which temperature and cerebral blood flow normalize rapidly (3, 4). Astonishingly, AGS suffer no ischemic injury during hibernation or reperfusion injury during an IBA. Hibernating ground squirrels are resistant to ischemic and reperfusion injuries in numerous models, including brain and heart tissues after cardiac arrest *in vivo* and hippocampal slice models derived from animals during an IBA (5-8). This resilience to reperfusion injury does not depend on temperature of the animal or season. In addition, AGS neural progenitor cells (NPCs) demonstrate resistance to oxygen and glucose deprivation *in vitro* (9). Together, these studies suggest that in addition to physiological adaptations, AGS possess cell autonomous genetic mechanisms that contribute to intrinsic tolerance to metabolic injury.

Proteomic and transcriptomic investigations have comprehensively catalogued the impact of season, torpor, and hibernation on cellular and metabolic pathways in several different tissues of hibernating ground squirrels, including brain (6, 10-16). Although the mechanisms underlying hibernating ground squirrel ischemia and hypothermia tolerance in the brain are not fully elucidated, studies suggest that post-translational modifications, regulation of cytoskeletal proteins, and upregulation of antioxidants play a prominent role (17-19). Recent studies employing unbiased next-generation sequencing and bioinformatics approaches have also highlighted the cytoprotective contributions of mitochondrial and lysosomal pathways in adapting to hypothermia and hypoxia in ground squirrel and marmot species (20, 21). In neurons differentiated from 13-lined ground squirrel (13LGS) induced pluripotent stem cells, Ou and colleagues found that hibernating ground squirrel microtubules retained stability upon exposure to hypothermia. The authors identified mitochondrial suppression of cold-induced reactive oxygen species (ROS) and preservation of lysosomal structure are key features of ground squirrel cytoprotection, and that pharmacological inhibition of ROS production or lysosomal proteases recapitulates the hypothermia-tolerant phenotype in human cells (21). Taken together, these studies have provided important insights into pathways mediating ischemia tolerance. However, these studies have not focused on specific proteins and their cytoprotective properties uniquely evolved in hibernating ground squirrels. As such, we know very little about mechanistic details underlying genetic contribution to intrinsic stress resilience in ground squirrels. Using a functional genetic screening approach combined with analyses of cell survival and mitochondrial phenotypes, we identified AGS transcripts imparting *in vitro* cytoprotection to various metabolic stressors. We further use CRISPR/Cas9 base editing (22) to determine functional importance of amino acid substitutions uniquely evolved in AGS, and identified AGS ATP5G1^L32^ as a mediator of key cytoprotective metabolic functions, suggesting potential for targeting this component of ATP synthase for neuroprotective treatments.

## Results

### AGS neural cells exhibit marked resistance to metabolic stressors associated with improvements in mitochondrial function and morphology

When growing under identical cell culture conditions, AGS and mouse NPCs exhibit similar morphology, growth rates and expression of Nestin and Ki67, markers for proliferating NPCs (Figure 1A and 1 Supplement). Although superficially indistinguishable, mouse and AGS NPCs demonstrate markedly different responses to metabolic stressors. When exposed to hypoxia (1% O_2_), hypothermia (31°C), rotenone (30 *µ*M), or FCCP (10 *µ*M), AGS NPCs exhibit profound resistance to cell death compared with mouse NPCs (Figure 1B), recapitulating resilient AGS phenotypes found in previous studies (5, 7-9). Moreover, measurement of *in vitro* oxygen consumption of AGS NPCs after sequential exposure to mitochondrial toxins demonstrates strikingly higher ‘spare respiratory capacity’ in response to FCCP (Figure 1C), indicating a greater metabolic reserve for stressors (23). Interestingly, these functional improvements in mitochondrial function were also mirrored by changes in mitochondrial dynamic organization following exposure to FCCP. At baseline, mouse and AGS cells had similar mitochondrial organization as evidenced by similar mean branch length and number of cells with fragmented mitochondria (Figure 1D,E). Following FCCP, an uncoupling agent that causes mitochondrial depolarization, mouse cells demonstrated large increases in mitochondrial fission with concurrent decreases in mean branch length. By contrast, AGS cells appeared largely resistant to mitochondrial fission induced by FCCP (Figure 1F,G). Together, these results demonstrate intrinsic differential cell survival and mitochondrial responses to metabolic stresses between mouse and AGS NPCs.

**Figure 1:**
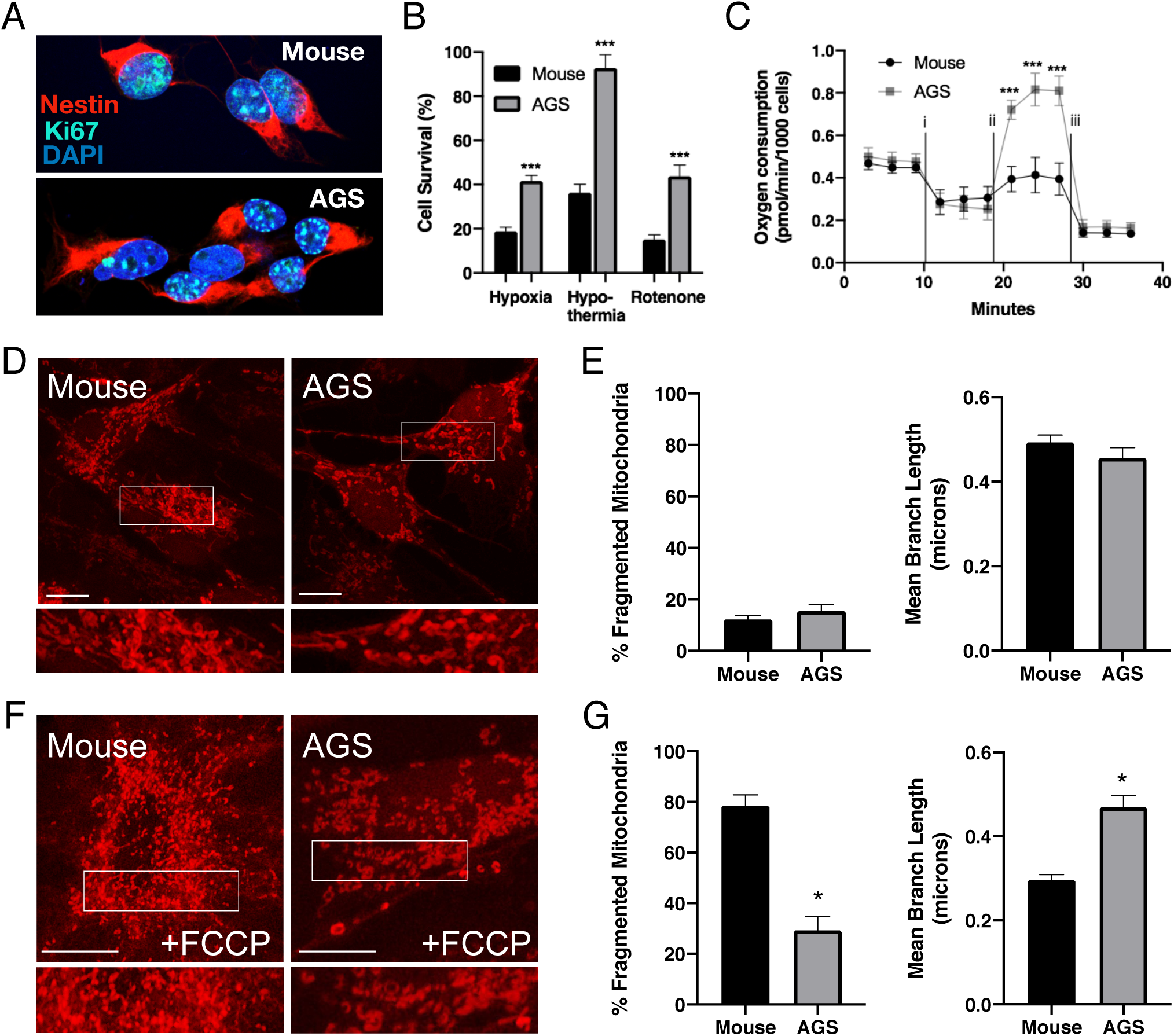
Phenotypic characteristics of Mouse and AGS NPCs. **(A.)** Confocal image of Mouse (top) and AGS (bottom) NPCs demonstrating similar morphology and expression of Nestin (red) and Ki-67 (teal) in nearly all cultured cells of both species. **(B.)** AGS NPCs exhibit increased cell survival when exposed to hypoxia (1%, 24hrs), hypothermia (31°C, 24hrs), or rotenone (10 μM, 16 hrs). Bar graphs represent the mean of 3 independent experiments with 3 replicates/condition. **(C.)** Seahorse XF analyzer assay of cultured mouse and AGS NPCs sequentially exposed to i) oligomycin (1μM), ii) FCCP (2μM), and iii) rotenone (0.5μM) showing enhanced FCCP-stimulated oxygen consumption (spare respiratory capacity). Data represents the mean of 3 independent experiments with 4-6 replicates/species. **(D and F.)** Representative confocal images of mouse (left) and AGS (right) NPCs expressing the mitochondrial marker mCherry-mito7 to demonstrate mitochondrial morphology at baseline (D.) and one hour following treatment with FCCP (F.). Scale bar represents 10 μm. **(E and G.)** Percent of mitochondria with fragmented morphology (left panel) and the mean branch length (right panel) of mitochondrial networks of NPCs expressing mCherry-mito7. Data obtained from 30 cells/species/condition. \**P*<0.05; \*\*\**P*<0.001.

### A cDNA library expression screen identifies AGS ATP5G1 as a cytoprotective factor

To identify cytoprotective genes expressed in AGS, we constructed a normalized cDNA expression library from AGS NPCs and introduced the library to mouse NPCs by nucleofection. Two days after AGS cDNA library nucleofection, we exposed cells to hypothermia (31°C) for 3 days, hypoxia (1%) for 2 days, or complex I inhibition (rotenone) for 3 days, respectively (Figure 2A and 2 Supplement). We then isolated plasmids from surviving cells, amplified cDNA insert sequences by PCR and used next-generation sequencing to identify a total of 378 putative cytoprotective genes, three of which (*Ags Atp5g1, Ags Manf*, and *Ags Calm1*) provided cytoprotection in all conditions (see Figure 2B and Supplemental Data File 1).

**Figure 2:**
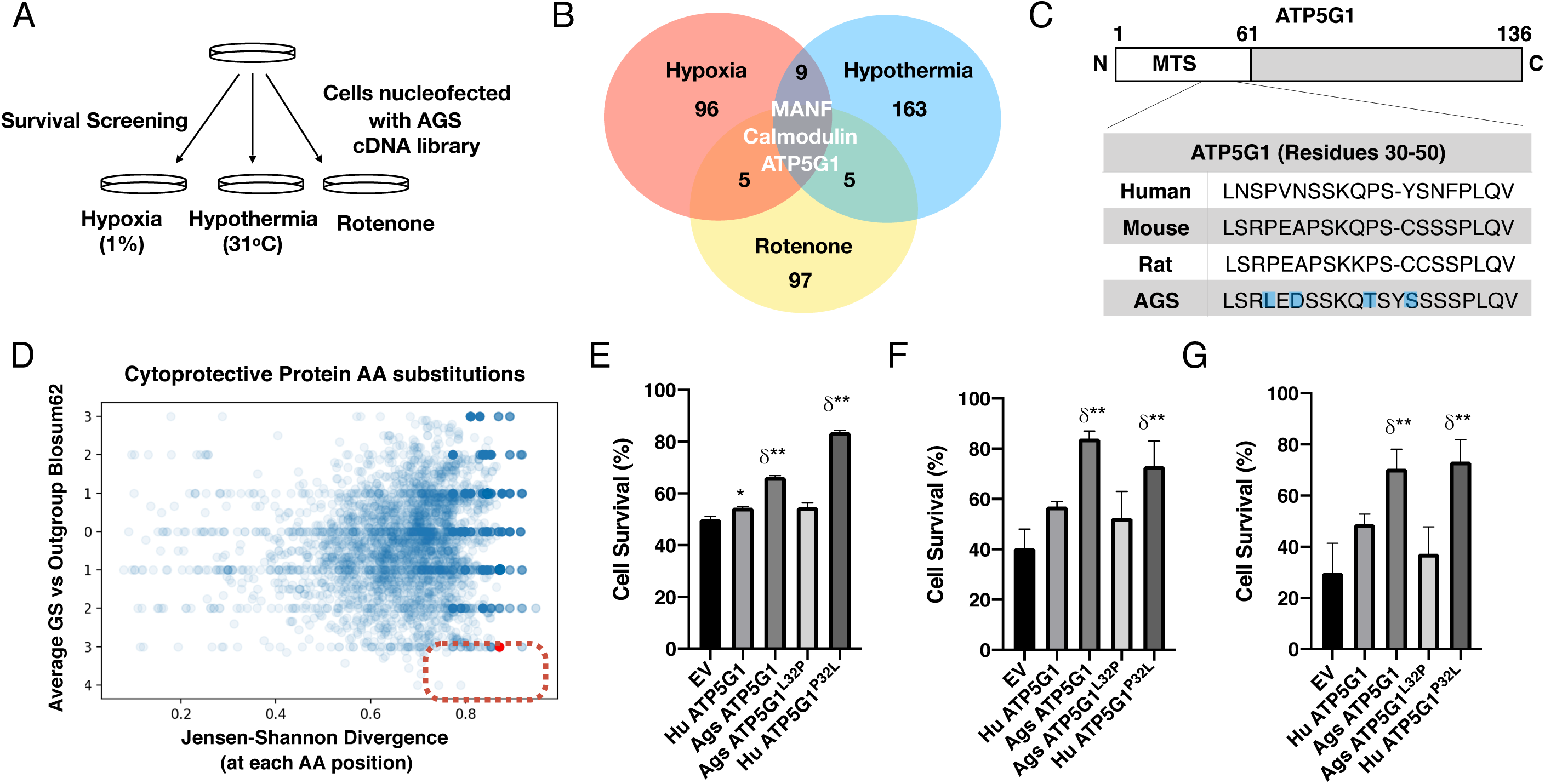
AGS cDNA library survival screen identifies AGS ATP5G1 as a cytoprotective factor. **(A.)** AGS NPC cDNA was introduced into mouse NPCs by nucleofection. Cells were screened for survival after exposure to hypoxia (1%, 48hrs), hypothermia (31°C, 72 hrs), or rotenone (20 μM, 48hrs) to identify AGS cytoprotective factors. **(B.)** Venn-diagram demonstrating the number of cytoprotective proteins identified by next-generation sequencing of plasmids isolated from cells surviving each condition of the cDNA library screen. **(C.)** Truncated sequence alignments demonstrating key GS AA substitutions (blue highlight) for ATP5G1, one of the three proteins imparting survival in all three screens. **(D.)** Ground squirrel-unique amino acid substitutions are plotted as a function of BLOSUM62 score and Jensen-Shannon Divergence (JSD) score. Ground squirrel unique AA substitutions with a higher probability of functional consequence are in the denoted red quadrant (high JSD values and low BLOSUM62 scores). The red dot represents the ATP5G1^L32P^ substitution; orange dots represent the other ATP5G1 substitutions. **(E-G.)** Mouse NPCs expressing human ATP5G1, AGS ATP5G1, AGS ATP5G1^L32P^, human ATP5G1 ^P32L^, or empty vector (EV) and exposed to 24 hours of 1% O_2_ (E.), 31°C (F.), or 20mM rotenone (G.). Cell death was determined by flow cytometry for propidium iodide and experiments are mean of 3 independent experiments with 3 replicates/genotype/condition, \**P*<0.05 or \*\**P*<0.01 vs EV; δ<0.05 vs human ATP5G1.

Since a portion of mouse NPCs survived metabolic stresses even without AGS cDNA library expression, we anticipated false positive hits and focused on the nuclear-encoded mitochondrial protein AGS ATP5G1, as it was one of three genes found to confer resilience across all three metabolic stress paradigms (Figure 2B). We hypothesized that resistance to mitochondrial insults may be related to uniquely evolved AGS amino acid substitutions in key proteins. There are three substitutions and two insertions/deletions in the AGS ATP5G1 N-terminal mitochondrial targeting sequence compared to the human and mouse sequences (Figure 2C and 2 Supplement C). ATP5G1 is one of three ATP5G isoforms making up the C subunit of ATP synthase, and is regulated distinctly from ATP5G2 of ATP5G3 (24-27). We hypothesized a subset of AGS cDNA library screened cytoprotective variants contain uniquely evolved amino acid substitutions. We identified uniquely evolved AGS proteins by comparing sequence alignments of the screened cytoprotective candidates for 2 species of ground squirrels (AGS and the 13LGS, Ictidomys tridecemlineatus) against 9 other reference species across mammalian subclasses. Jensen Shannon Divergence (JSD) score, and average ground squirrel-versus-other mammalian block substitution matrix (BLOSUM)-62 scores were calculated for each unique residue, as described previously (28). JSD was used to capture sequence conservation and difference from the background amino acid distribution. High JSD and low BLOSUM62 scores indicate chemically significant amino acid substitutions, and thus may indicate potentially important functional AGS adaptations. The leucine-32 of AGS ATP5G1 found in place of the highly conserved proline is unique to hibernating ground squirrel, and on conservation analysis scored among the highest of all AGS-unique amino acid substitutions in screened cytoprotective proteins (28; Figure 2D and Supplemental Data File 2).

The N-terminal region of ATP5G proteins can undergo cleavage, but also modulate mitochondrial function directly, though the specifics of the mechanism are unknown (29). Although the three C subunit proteins are identical in sequence, they cannot functionally substitute for one another and are all required to constitute a fully functional C subunit (29, 30). To determine the relative levels of ATP5G1, −2, and −3 in mouse and AGS NPCs, we performed qRT-PCR analysis with species and transcript-specific primers. We found that in both mouse and AGS NPCs, expression of *Atp5g3* or *Atp5g2* is greater than that of *Atp5g1*, consistent with prior reports in human and mouse tissues (25, 29). However, the relative abundance of the *Atp5g1* isoform is elevated nearly two-fold in AGS NPCs (Figure 2 Supplement D). Interestingly, the relative abundance of the mature ATP5G (subunit C) protein or complex V activity, is not different in mouse or AGS cells (Figure 2 Supplement E), suggesting that the beneficial effects of ATP5G1 could be mediated by interaction with other proteins.

Overexpression of the AGS variant of *ATP5G1* in mouse NPCs confers cytoprotection in cells exposed to hypoxia, hypothermia, or rotenone. We found that this protective response is not present in NPCs overexpressing ATP5G1^*L32P*^. Similarly, overexpression of the human ATP5G1^*P32L*^, which mimics the wild-type AGS ATP5G1 variant, leads to enhanced cytoprotection in these conditions of metabolic stress compared to human ATP5G1 (Figure 2E-G). The AGS ATP5G1 substitutions did not alter the mitochondrial localization of ATP5G1 (Figure 3 Supplement A-C), and importantly, recapitulated key features of the AGS resilient mitochondrial phenotype, including increasing spare respiratory capacity and reducing mitochondrial fission with reduced fragmentation and increased branch length of mitochondria (Figure 3A-D). Of note, the other identified ground squirrel amino acid substitutions (N34D, T39P) did not demonstrate a loss of the protective effect on survival when overexpressed in mouse NPCs exposed to hypoxia, hypothermia, or rotenone (Figure 3 Supplement D-F).

**Figure 3:**
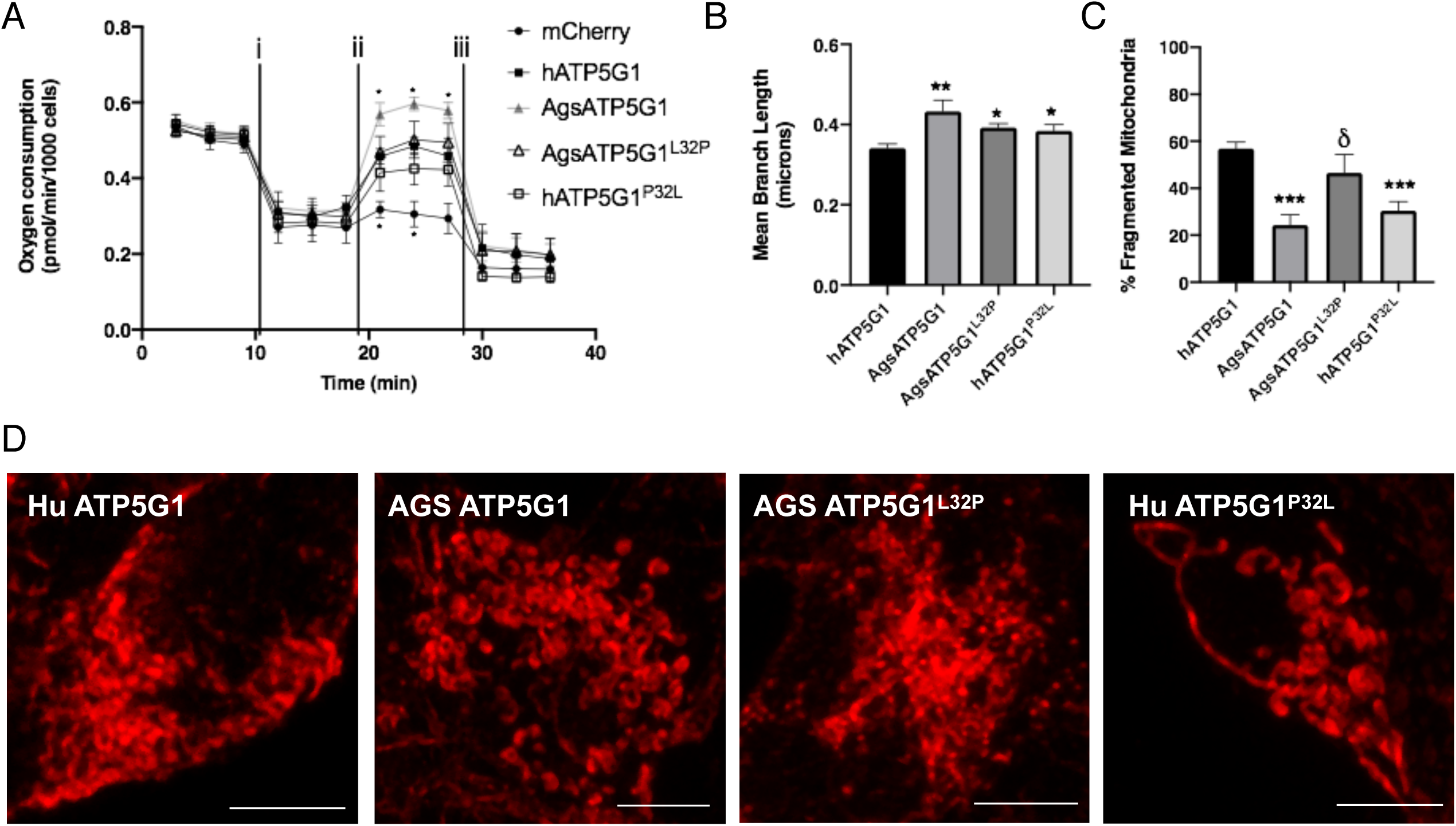
Overexpression of AGS ATP5G1 in mouse NPCs recapitulates AGS metabolic phenotypes, which is dependent on the uniquely evolved leucine-32. **(A.)** Seahorse XF analyzer assay of cultured mouse NPCs expressing human ATP5G1, AGS ATP5G1, AGS ATP5G1^L32P^, human ATP5G1^P32L^, or empty vector and sequentially exposed to i) oligomycin (μM), ii) FCCP (2μM), and iii) rotenone (0.5μM) showing increased FCCP-stimulated oxygen consumption (spare respiratory capacity) with AGS ATP5G1. Substitution of the AGS leucine-32 results in reduced spare respiratory capacity. **(B-D.)** Representative confocal images (D.) of mitochondrial networks in mouse NPCs expressing human, AGS, and mutant forms of mCherry-ATP5G1 one hour following treatment with FCCP. Percent of mitochondria with fragmented morphology (B.) and the mean branch length (C.) of mitochondrial networks of ATP5G1 overexpressing mouse NPCs. Data obtained from 30 cells/condition. \**P*<0.05; \*\**P*<0.01 vs. human ATP5G1; δ<0.05 vs AGS ATP5G1. Scale bar in (D.) represents 5 um.

### Knock-in of AGS ATP5G1^L32P^ alters the resilient phenotype of AGS cells

Using the previously described adenine base editor (ABEmax; 22), we successfully generated AGS cell lines homozygous for ATP5G1^*L32P*^ by introducing a cytosine-to-thymine substitution in the (-) strand of *Ags Atp5g1* (Figure 4A,B). We isolated three clonal AGS NPC lines harboring the desired knock-in mutation (ABE KI) and two clonal lines that underwent editing and remained homozygous for the wild-type ATP5G1 gene (ABE WT). Compared to ABEmax-treated AGS cells without successful knock-in (Figure 4 Supplement A), ABE KI cell lines did not demonstrate differences in ATP5G mRNA, protein expression, or complex V activity (Figure 4 Supplement B-E). In spite of stable ATP5G protein and complex V activity, knock-in of the L32P residue resulted in reduced survival following exposure to hypoxia, hypothermia, or rotenone (Figure 4C), again indicating that the cytoprotective effects of ATP5G1 may be mediated by interactions with other proteins. In addition, we found the ABE KI AGS NPCs exhibited marked reduction in ‘spare respiratory capacity’ and altered mitochondrial dynamics in response to FCCP treatment (Figure 4D-G). Taken together, these results support a role for the AGS Leucine-32 substitution in the N-terminus of ATP5G1 in mediating cytoprotection against metabolic stresses by modulating mitochondrial function.

**Figure 4:**
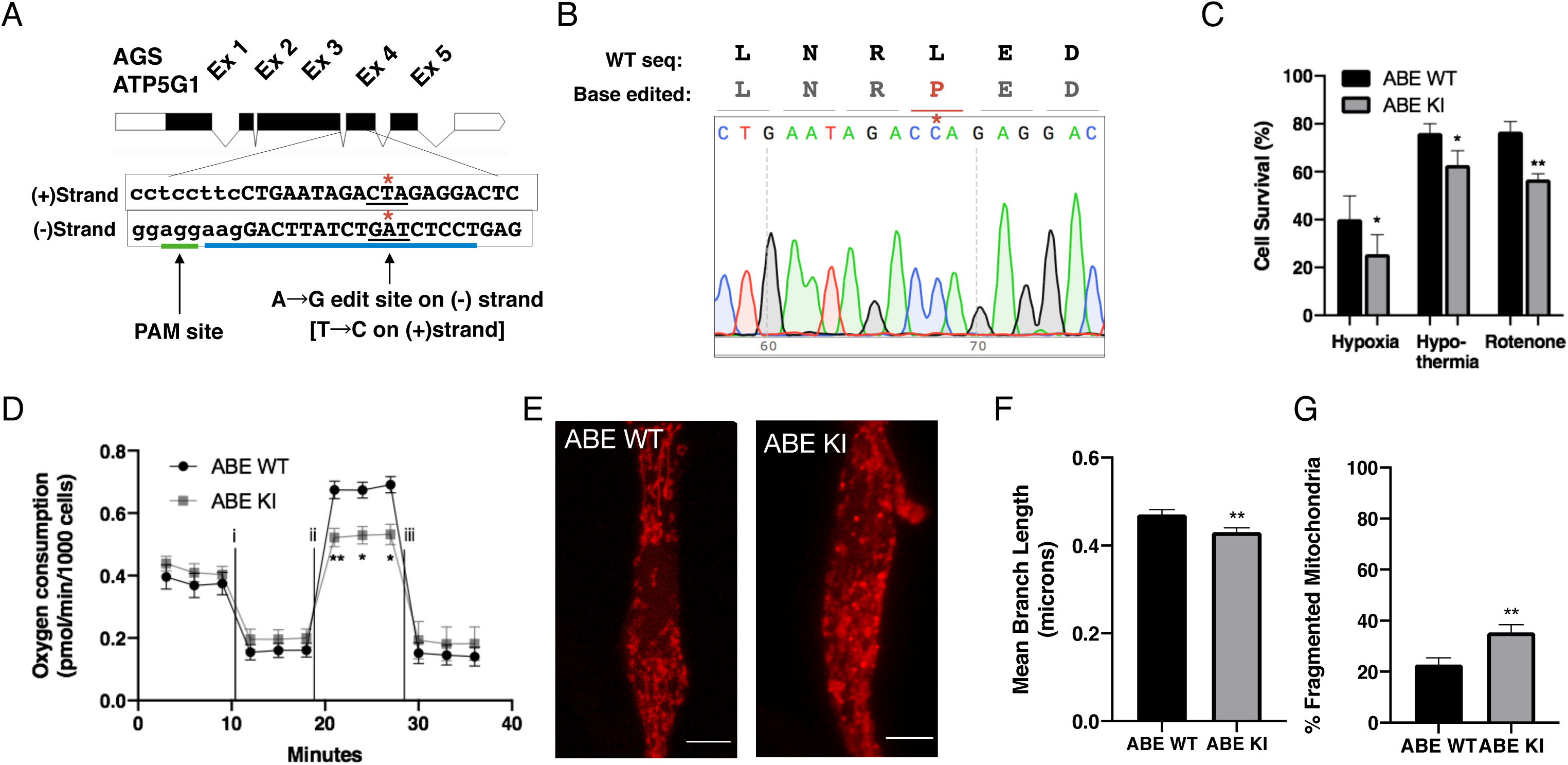
CRISPR base editing to generate ATP5G1^L32P^ AGS NPCs results in a partial loss of AGS metabolic resilient phenotypes. **(A.)** AGS ATP5G1 CRISPR base editing strategy. To create AGS cells with the human amino acid substitution at leucine-32, AGS cells transiently expressing ABEmax were nucleofected with an sgRNA (blue underline) directed towards a PAM site (green underline) on the (-) strand to target conversion of adenine to guanine, which on the (+) strand is a cytosine to thymine (*). **(B.)** Sequencing data from a successfully edited clonal AGS cell line demonstrating the cytosine to thymine base edit resulting in the desired leucine to proline knock-in cell line. **(C.)** AGS ATP5G1^L32P^ (ABE KI) NPCs exhibit decreased cell survival compared to unedited AGS cells (ABE WT) when exposed to hypoxia (1%, 24hrs), hypothermia (31°C, 72hrs), or rotenone (10 μM, 16 hrs). Bar graphs represent the mean of 3 independent experiments with 3 replicates/condition. **(D.)** Seahorse XF analyzer assay of cultured ABE KI and WT cells sequentially exposed to i) oligomycin (1uM), ii) FCCP (2uM), and iii) rotenone (0.5uM) showing enhanced FCCP-stimulated oxygen consumption (spare respiratory capacity). Data represents the mean of 3 independent experiments with 4-6 replicates/species. **(E.)** Representative confocal images of ABE WT (left) and ABE KI (right) NPCs expressing the mitochondrial marker mCherry-mito7 to demonstrate mitochondrial morphology one hour following treatment with FCCP. Scale bar represents 5 um. **(F and G.)** Percent of mitochondria with fragmented morphology (F.) and the mean branch length (G) of mitochondrial networks of ABE WT and KI NPCs expressing mCherry-mito7. Data obtained from 50-60 cells/genotype. \**P*<0.05; \*\**P*<0.01.

## Discussion

Previous studies have indicated that organisms capable of hibernation evolved a myriad of physiological and cellular mechanisms to survive under metabolic stress conditions during the hibernation process (20, 21, 31). However, we still know little about how specific protein-coding genetic variants evolved in the AGS genome causally contribute to mechanisms mediating intrinsic cytoprotective functions. We show that *ex vivo* cultured AGS NPCs demonstrate remarkable intrinsic resilience to hypoxia, hypothermia, and other metabolic stressors. Additionally, using an unbiased cDNA expression screening and bioinformatic strategy, we identified numerous AGS transcripts and uniquely evolved AGS amino acid substitutions potentially contributing to cytoprotection. We focused on discerning the protective effect of AGS ATP5G1, a nuclear-encoded mitochondrial protein, given that it was one of only three genes identified in all three metabolic stress paradigms and the prominent mitochondrial resilience phenotype of AGS NPCs. We hypothesize that analogous to amino acid substitutions in several human proteins providing adaptive benefits (32-35), substitutions in AGS ATP5G1 may underlie AGS adaptive mechanisms contributing to its robust cytoprotective phenotype. Utilizing dCas9 ABE technology, we validated a unique *Ags* ATP5G1^*L32*^ amino acid substitution in the mitochondrial targeting sequence of ATP5G1 that leads to improvements in mitochondrial physiologic parameters. Thus, this study is among the first to our knowledge to use CRISPR base editing in a non-model organism, and demonstrates a potential role for modulating ATP5G1 to enhance cytoprotection in ischemic diseases. Furthermore, the novel mitochondrial functional and morphological phenotypes established here can serve as a robust *in vitro* paradigm to investigate the numerous other identified gene perturbations potentially mediating the AGS resilient phenotype with implications for novel neuroprotective treatments as well as promoting survival of neural stem cell grafts (36).

Mitochondrial metabolic dysfunction is central to ischemia and reperfusion injury. Physiologic, transcriptomic, and proteomic studies have highlighted the importance of ketone and fatty acid metabolism in hibernating states (37, 38) as well as pointed to a role for specific post-translational protein modifications in the differential regulation of metabolic pathways in hibernation (10, 39, 40). We expanded this body of knowledge, by identifying altered mitochondrial dynamics and enhanced spare respiratory capacity in cells of hibernating species as potentially adaptive cellular mechanisms in hibernating animals. This mitochondrial phenotype is likely responsible for the broad resilience of AGS cells against a wide range of metabolic stressors. The downstream mediators of this substitution’s beneficial effect remain to be determined, but do not appear to involve regulation of the mature ATP5G protein abundance or complex V activity.

Spare respiratory capacity, as measured by FCCP-stimulated oxygen consumption, represents a marker for cellular metabolic reserves (23) and is thought to be determined by the oxidative phosphorylation machinery (41, 42). AGS demonstrate marked elevations in spare respiratory capacity compared to mouse cells. While this is likely the result of AGS adaptations in numerous proteins, the importance of the ATP5G1 variant is highlighted by our experimental evidence demonstrating improvement in spare respiratory capacity in mouse NPCs over-expressing AGS ATP5G1 variants and decreased spare respiratory capacity in AGS NPCs with ATP5G1 L32P knock-in. A critical role for ATP5G1 in cellular energetics is also supported by recent work uncovering ATP5G1 as one of the major effectors of the transcription factor, BCL6, in regulating adipose tissue energetics as well as maintaining thermogenesis in response to hypothermia (43, 44).

Mitochondrial fission and fusion are regulated by cellular metabolic state and a host of regulatory proteins, many of which have been implicated in cell survival response to stresses (45). While metabolic stresses often lead to mitochondrial fission followed by apoptosis, mitochondrial fusion and resistance to fission in response to stress are anti-apoptotic (46, 47). Fusion is hypothesized to allow for complementation of damaged and dysfunctional mitochondria, and in states of metabolic stress, hyperfusion of mitochondria helps maintain mitochondrial membrane potential and cell viability (48). The increase in fusion and improvement in cell survival in mouse NPCs over-expressing AGS ATP5G1 and loss of resilient metabolic phenotypes in AGS cells carrying ATP5G1^*L32P*^ underscore the importance of this protein in altering mitochondrial morphologic response to stresses and increasing the metabolic oxidative capacity of cells.

In mammals, all of the approximately 1000 nuclear-encoded mitochondrial proteins contain a unique mitochondrial targeting sequence (MTS) providing a high degree of specificity in regulating mitochondrial import and sorting. These mitochondrial targeting and processing functions are regulated by the highly conserved mitochondrial membrane translocating protein complexes (TOM and TIM) and MTS cleaving proteins, mitochondrial processing peptide (MPP) and mitochondrial intermediate peptide (MIP). We did not find evidence that the ATP5G1 MTS sequence differences between AGS and mouse or ABE WT and KI affected the mitochondrial localization or expression levels of mature ATP5G protein, suggesting that the phenotypic differences observed are not related to mitochondrial protein import or processing. Furthermore, the putative ATP5G1 ten amino acid sequential MPP/MIP cleavage site (xRx↓(F/L/I)xx(S/T/G)xxxx↓; 49) is also completely conserved in human, mouse, and AGS (see Figure 2 Supplement C). In addition, the MTS of proteins can vary greatly in their length, and many have speculated that in addition to organelle targeting the MTS also play important regulatory roles (50). The cleaved ATP5G1 N-terminal mitochondrial targeting sequence has been described to modulate mitochondrial function distinct from the functionally active C-terminal protein (29). However, additional experiments will be required to understand how the AGS substitutions in ATP5G1 mediate the cytoprotective effect. Interestingly, although increased abundance of the *Atp5g1* transcript in AGS neural cells may also contribute to the altered mitochondrial function in brown adipose tissue (27), the ABE KI cells did not demonstrate a difference in ATP5G1 transcript abundance suggesting that ATP5G1 transcriptional isoform abundance does not underlie ATP5G1-related cytoprotection.

Further unraveling of the mechanisms underlying AGS mitochondrial and cellular resilience to metabolic injuries holds the hope of identifying novel cytoprotective strategies that will lead to improved cellular engineering strategies and treatments for ischemic diseases. Systematic experimental investigation of the additional amino acid substitutions identified in AGS will provide important insights into the pathways underlying intrinsic ischemic tolerance. The use of CRISPR gene editing technologies coupled with detailed phenotypic analysis is a unique and powerful approach to evaluate causal roles of genetic variants in conferring phenotypic traits. Identification of such causal variants for stress resilience in AGS may help develop pharmacological, gene therapy, or CRISPR/genome editing-based therapeutic strategies to treat human ischemic disorders, including stroke and heart attack.

## Materials and Methods

### Cell culture

AGS NPCs (Neuronascent, Gaithersburg, MD, USA) and mouse NPCs (gift of Song lab, Baltimore, MD) have been previously described (9, 51). They were grown under standard conditions at 37°C and 5% CO_2_ with NeuroCult basal media (STEMCELL, Vancouver, BC, CA) with EGF (50 ng/ml, PeproTech, Inc., Rocky Hill, NJ, USA), FGF (100 ng/ml, PeproTech, Inc.), heparin (0.002%), and proliferation supplements (STEMCELL). Early passage cultures (P2) were expanded and frozen and thawed in batches for use in experiments. These cultures contain cells ubiquitously expressing the NPC marker, Nestin, and the proliferation marker, Ki-67 (Figure S1). For *in vitro* modeling of metabolic stress, cells were exposed to either: i) 1% hypoxia in a specialized incubator (Nuaire, Plymouth, MN, USA) saturated with Nitrogen/5% CO_2_; ii) hypothermia in standard incubators maintained at lower temperatures; and iii) complex I inhibition with the addition of rotenone to cell media.

### DNA constructs and lentiviral transfection

The pHAGE-ATP5G plasmids were generated by direct PCR and PCR fusions; and the point mutation plasmids generated using Q5 site-directed mutagenesis (New England Biolabs, Beverly, MA, USA). For lentiviral transfection, the plasmids with packaging plasmids were co-transfected into HEK293FT (with a ratio of 2:1.5:1.5) using Turbofect reagent (Thermo Fisher Scientific Inc., Waltham, MA, USA) according to the manufacturer’s instructions. Lentivirus-containing medium was filtered from the post-transfection supernatant and used for transduction of HEK293T cells or mouse NPCs. All lentivirus-infected cells were cultured in the medium containing Polybrene (4 μg/ml; Sigma Aldrich, St. Louis, MO, USA) for 8 hours before changing media. Forty-eight hours after transduction, the cells were selected with 10 *µ*g/ml Blastidicin S (Thermo Fisher Scientific Inc.).

### Generation of CRISPR Base-edited AGS cells

ATP5G1^L32P^ NPCs were generated using the dCas9 base editor, ABEmax (Addgene #112095), as previously described. Briefly, a synthetic sgRNA (TCCTCTAGTCTATTCAGGAA) was selected by manual inspection of the Ags *Atp5g1* sequence for a PAM (NGG) site near the desired edit on the (-) strand of the gene. AGS NPCs were nucleofected (Amaxa 4D, program DS113) in P3 solution (Lonza, Alpharetta, GA, USA) containing pCMV ABEmax (500 ng/200,000 cells). Following a 48 hour recovery period, the same cells were nucleofected with the synthetic sgRNA sequence above (100 pmol, Synthego, Menlo Park, CA, USA). Cells were expanded and then clonally plated. Clones were screened by PCR as the desired base edit also introduced a new BfaI restriction enzyme cutting site. Sanger sequencing was used to confirm the two WT and three KI clone sequences utilized. Potential off-target effects of CRISPR/Cas9 cleavage were analyzed by Sanger sequencing of the top 5 predicted off-target genomic locations [https://mit.crispr.edu], which demonstrated a lack of indels for all clones used in subsequent analysis.

### Cell death assay

Mouse and AGS cells were plated in 24 or 96-well plates and grown to 70% confluence. Cells were exposed to metabolic stress paradigms as above, and detached and floating cells collected by centrifugation and washed with 1 ml PBS. The collected cells were resuspended with 200 μl PBS with addition of 0.2 μl Sytox blue (1 *µ*M; Thermo Fisher Scientific) or propidium iodide (concentration) for an additional 5 min. The fluorescence intensity was measured for individual cells using automated cytometry (Arthur, Nanoentek, Waltham, MA, USA) or flow cytometry (BD Biosciences, San Jose, CA, USA) within 20 min of staining, and the percentage of cell death quantified using the FlowJo software.

### cDNA Library generation, screening, and identification of AGS amino acid substitutions

RNA was isolated from AGS NSC/NPC cells grown under standard conditions. A normalized cDNA library was generated by a commercial research partner (Bio S&T, Montreal, QC, Canada) from RNA extracted from AGS NPCs. Library quality and normalization is shown in Figure 2 Supplement A and B). For library screening, plates containing 1 × 10^7^ mouse NPCs cells were grown in triplicate and nucleofected with 200,000 clones each. Plates were exposed to one of three metabolic stress conditions (hypoxia, hypothermia, or rotenone treatment) for 48-96 hours. Following this treatment, plasmid DNA was purified from surviving cells and PCR-amplified AGS cDNA inserts subjected to next-generation sequencing. Resulting fastq files were trimmed (Trim Galore!) and mapped to the Ictidomys Tridecemlineatus genome (SpeTri2.0) using HISAT2. Mapped reads were subjected a custom pipeline for analyzing amino acid substitutions. Briefly, protein sequences of mapped genes were queried by gene symbol and downloaded from OrthoDBv10 (52) for 10 species (13LGS, Mus musculus, Rattus norvegicus, Sorex araneus, Pongo abelii, Homo sapiens, Equus caballus, Bos taurus, Oryctolagus cuniculus, Sus scrofa). OrthoDB data was filtered by matching records against accepted GeneCards aliases for each gene. Multiple records per species were resolved using maximum percent identity against the accepted human, mouse, and 13LGS sequences, such that only one record per species was used for alignment. AGS protein sequences were downloaded from the Entrez Protein database. Multiple AGS isoforms were resolved by best identity match to the OrthoDB sequence data. The final protein sequence set was aligned with KAlign 2.04 (53). From aligned sequences, GS-specific residue substitutions were defined as amino acid variants present in 13LGS and AGS sequences and present in no other included species. For each GS-specific residue, sequence weights, JSD, and average GS-versus-outgroup BLOSUM62 scores were calculated as described previously (28). BLOSUM62 scores were used instead of point-accepted mutation scores in order to prioritize protein sequence changes with higher probability of potential chemical and functional difference. JSD was used to capture sequence conservation and difference from the background amino acid distribution. BLOSUM62 scores were calculated for GS residues against all other mammalian species sequences and averaged to give GS vs Outgroup BLOSUM62. For the entire screened cytoprotective protein dataset, JSD and BLOSUM62 score were plotted for individual genes of interest against the remaining dataset.

### Analysis of in vitro mitochondrial respiration

Analysis of mitochondrial respiratory potential was performed using a flux analyzer (Seahorse XF^e^96 Extracellular Flux Analyzer; Seahorse Bioscience, North Billerica, MA, USA) with a Seahorse XF Cell Mito Stress Test Kit according to the manufacturer’s instructions. Basal respiration and ATP production were calculated to evaluate mitochondrial respiratory function according to the manufacturer’s instructions. After the measurement, cells were harvested to count the cell number, and each plotted value was normalized relative to the number of cells used. Briefly, NPCs were seeded (25,000 cells/well) into each well of XF^e^96 cell culture plates and were maintained in standard culture media. After 2-3 days in culture, cells were equilibrated in unbuffered XF^e^ assay medium (Seahorse Bioscience) supplemented with glucose (4.5 g/L), sodium pyruvate (25 mg/L) and transferred to a non-CO_2_ incubator for 1 h before measurement. Oxygen consumption rate (OCR) was measured with sequential injections of 1-2 *μ*M oligomycin, 2-4 *μ*M FCCP and each 0.5 *μ*M of rotenone/antimycin A.

### Analysis of mitochondrial respiratory chain complex activity

Analysis of mitochondrial respiratory chain complex V activity was measured with Complex V Mitocheck kit (Cayman Chemical, Ann Arbor, MI, USA). Mitochondrial extracts (50 μg) were obtained as previously described (54) and used to measure time-dependent absorbance alterations on a multi-well plate reader (SprectraMax, Molecular Devices, San Jose, CA, USA).

### Mitochondrial Dynamic Morphology Assessment

Mitochondrial morphology and fission/fusion is assessed in mouse and AGS NPCs nucleofected with mCherry-Cox8 as a mitochondrial marker and grown on glass coverslips in standard media. Cells are allowed to recover for 48 hours and then fixed with paraformaldehyde (4%) one hour following treatment with FCCP (1 uM) or DMSO. High magnification images of cells are captured by confocal microscopy and mitochondrial morphological characteristics were assessed with the Mitochondrial Network Analysis (MiNA) toolset in J-image as previously described (55, 56). Briefly, the plugin converts confocal images to binary pixel features and analyzes the spatial relationship between pixels. The parameters analyzed are: (i) individual mitochondrial structures; ii) networked mitochondrial; and iii) the average of length of rods/branches. Twenty randomly chosen fields containing 30-50 cells were used to quantify the morphological pattern and network branch lengths of mitochondria. We classify the mitochondrial morphology as fragmented when the appearance is completely dotted with branch lengths < 1.8 μm.

### Immunoblot analysis

Laemmli loading buffer (Bio-Rad Lab, Hercules, CA, USA) plus 5% β-mercaptoethanol was added to protein extracts from cell pellets before heating at 95 °C for 5 min. Around 30 μg of whole cell protein lysate samples were separated on mini-PROTEIN GTX precast gels, and transferred to nitrocellulose membranes (Bio-rad). After blocking in TBS containing 5% non-fat milk and 0.1% Tween-20, the membranes were incubated overnight with primary antibodies diluted in TBST (Tris buffered saline with Tween-20; Genesee Scientific) with 5% milk at 4 °C, followed by incubation with secondary antibodies at room temperature for 1 h. Immunoreactivity was visualized by the ECL chemiluminescence system (Bio-rad) on standard film. The antibodies were ATP Synthase C subunit (ab-181243, 1:1000, Abcam, Cambridge, MA), HSP90 (sc-69703, 1:1000, Santa Cruz Biotechnology, Dallas, TX, USA).

### Immunofluorescence and confocal microscopy

For immunocytochemistry of mammalian cells, AGS and mouse NSC/NPC cells were seeded on laminin-coated coverslips (Neuvitro, Vancouver, WA, USA) within 24-well plates. The cells were fixed with 4% paraformaldehyde in PBS, washed with PBS, and permeabilized with 0.02% Triton X-100 in PBS for 10 min. Blocking was done with 5% BSA in PBS for 1 h, followed by incubation with antibodies against Nestin (MAB2736, 1:50, R&D Systems, Cambridge, MA, USA) or Ki-67 (NB600-1252, 1:500, Novus Biologicals, Littleton, CO, USA) in blocking buffer overnight at 4 °C. The Nestin antibody was detected using goat anti-mouse AlexaFluor 488 or 647 (1:1000; Jackson ImmunoResearch Laboratories Inc., West Grove, PA, USA) and the Ki-67 antibody was detected using AlexaFluor 488 goat anti-rabbit (1:1000; Jackson Immunoresearch) or Cy3-conjuated donkey anti-rabbit (1:500; EMD Millipore, Burlington, MA, USA) in blocking buffer. Cells were washed with PBS after primary and secondary antibody staining. Stained cells were overlaid with Fluoroshield mounting medium with DAPI (Abcam) to label nucleus. Fluorescence microscopy was performed with a Leica confocal microscope using the following fluorescence filters: DAPI (405 nm excitation); Cy3 (551 nm excitation); AlexaFluor 647 (651 nm excitation); and GFP/AlexaFluor 488 (488 nm excitation). For comparison across conditions, identical light-exposure levels were used.

### Quantitative RT-PCR

RNA was extracted from approximately 200,000 mouse or AGS NPCs per condition according to manufacturer instructions (Quick-RNA MiniPrep kit; Irvine, CA, USA). Total RNA was reverse transcribed into cDNA (Bimake, Houston, TX, USA), and real-time PCR was performed (LightCycler96, Roche, Basel, CHE) with SYBR Green (Thermo Fisher Scientific) as a dsDNA-specific binding dye. Quantitative RT-PCR conditions were 95°C for denaturation, followed by 45 cycles of: 10 s at 95°C, 10 s at 60°C, and 20 s at 72°C. Species-specific primers for each transcript were used (for list see Supplemental Table 1). Melting curve analysis was performed after the final cycle to examine the specificity of primers in each reaction. Relative abundance of each *Atp5g* isoform as a fraction of total *Atp5g* was calculated by ΔΔCT method and normalized to *Rpl27*.

### Statistical Analysis

Data were analyzed using GraphPad Prism Software (Graphpad, San Diego, CA) and presented as means ± S.E. unless otherwise specified, with *P-*values calculated by two-tailed unpaired Student’s *t*-tests or two-way ANOVA (comparisons across more than two groups) adjusted with the Bonferroni’s correction. No randomization or blinding was used and no power calculations were done to detect a pre-specified effect size.

### Data availability

The data that support the findings of this study are available from the corresponding author upon reasonable request.

## Acknowledgments

NSS and MB receive support from the American Heart Association, 18CDA34030443 and 19POST34381071, respectively. DKM receives support from National Institutes of Health grant R01GM117461, Pew Scholar Award, Curci Faculty Scholar Award from the Innovative Genomics Institute, and Packard Fellowship in Science and Engineering. The authors acknowledge use of sponsored core facilities including the UCSF Laboratory for Cell Analysis (P30CA082103).

## Author Contributions

NSS, MB, ELM, DKM conceived of the work, designed the experiments, and interpreted the data. NSS, MB, ELM, SL, and KRC performed experiments and acquired and analyzed the data. NSS drafted and along with all other authors revised the manuscript. DKM conceptualized and supervised the project.

---

**Figure 1 Supplement:**
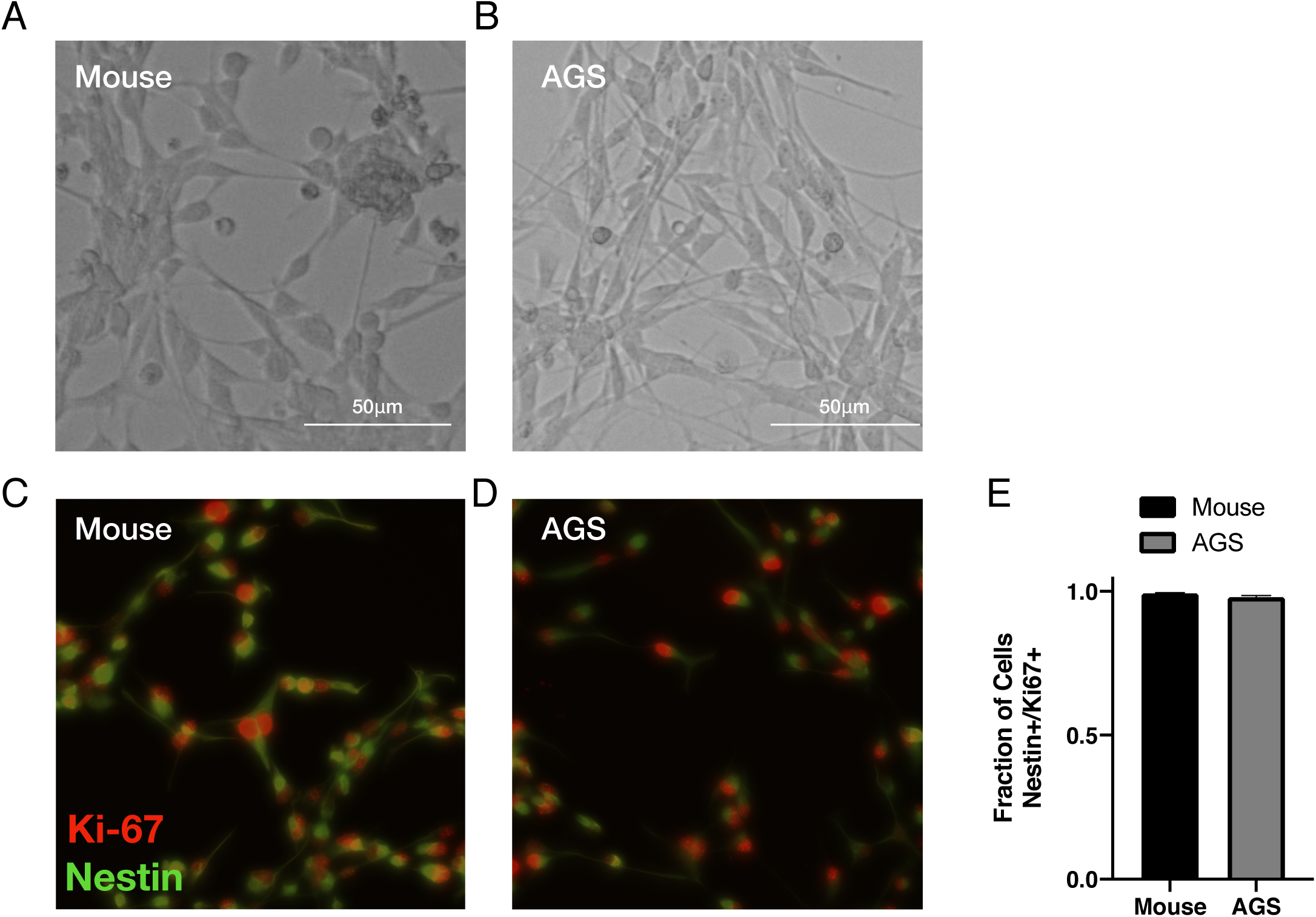
Mouse and AGS NPCs express Nestin and Ki-67. **(A-B.)** Representative live brightfield microscopy image of mouse and AGS NPCs. **(C-D.)** Representative fluorescent microscopy images of fixed mouse and AGS NPCs immunolabeled with Nestin (green) and Ki-67 (red). **(E.)** Quantitative assessment of mouse and AGS NPCs demonstrates that nearly all cells express both Nestin and Ki-67. Cell counts from 25 microscopic fields/cell line.

**Figure 2 Supplement:**
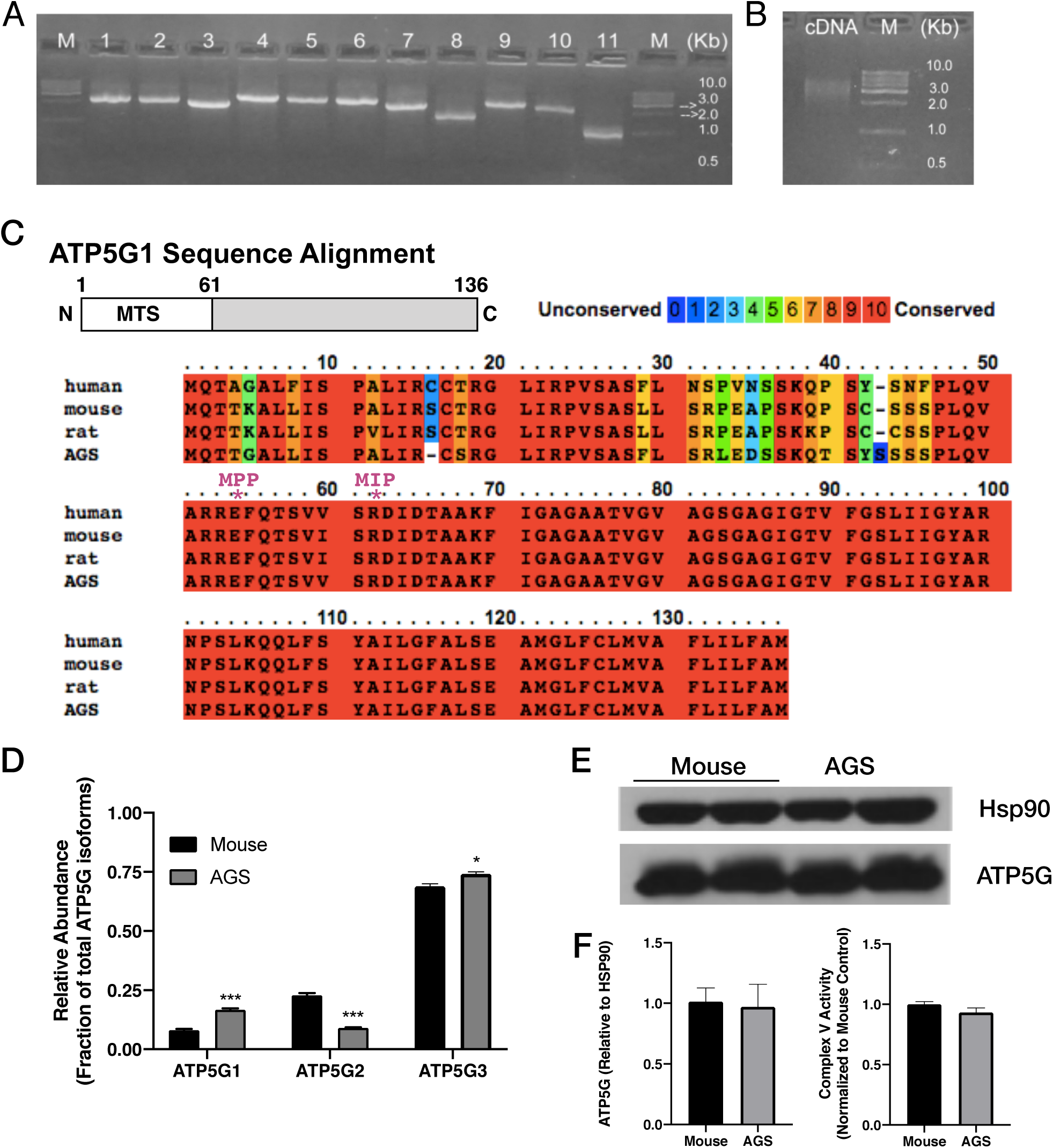
cDNA library construction and ATP5G expression in mouse and AGS NPCs. **(A.)** Agarose gel of 11 randomly chosen AGS cDNA library clones prior to amplification demonstrating an average insert size of 2.4 kB. **(B.)** Agarose gel of amplified and normalized AGS cDNA library. **(C.)** ATP5G1 sequence alignment in human, mouse, AGS, and rat demonstrating variability in the mitochondrial targeting sequence (alignment visualized using PRALINE, http://www.ibi.vu.nl/programs/pralinewww/). The **\*** indicates putative mitochondrial processing peptide (MPP) and mitochondrial intermediate peptide (MIP) cleavage sites. **(D.)** qRT-PCR for ATP5G1, ATP5G2, and ATP5G3 demonstrating increased relative abundance of ATP5G1 in AGS NPCs. Data from 4 independent experiments performed in triplicate. \**P*<0.05\*\*\**P*<0.001. **(E.)** Representative western blot images demonstrating the relative abundance of ATP5G protein isoforms is similar in mouse and AGS. **(F.)** Quantification of western blots from 4 independent experiments with 2-3 replicates each. **(G.)** Complex V activity measured in mitochondrial extracts from mouse and AGS. Data from 3 independent experiments with 3 replicates each.

**Figure 3 Supplement:**
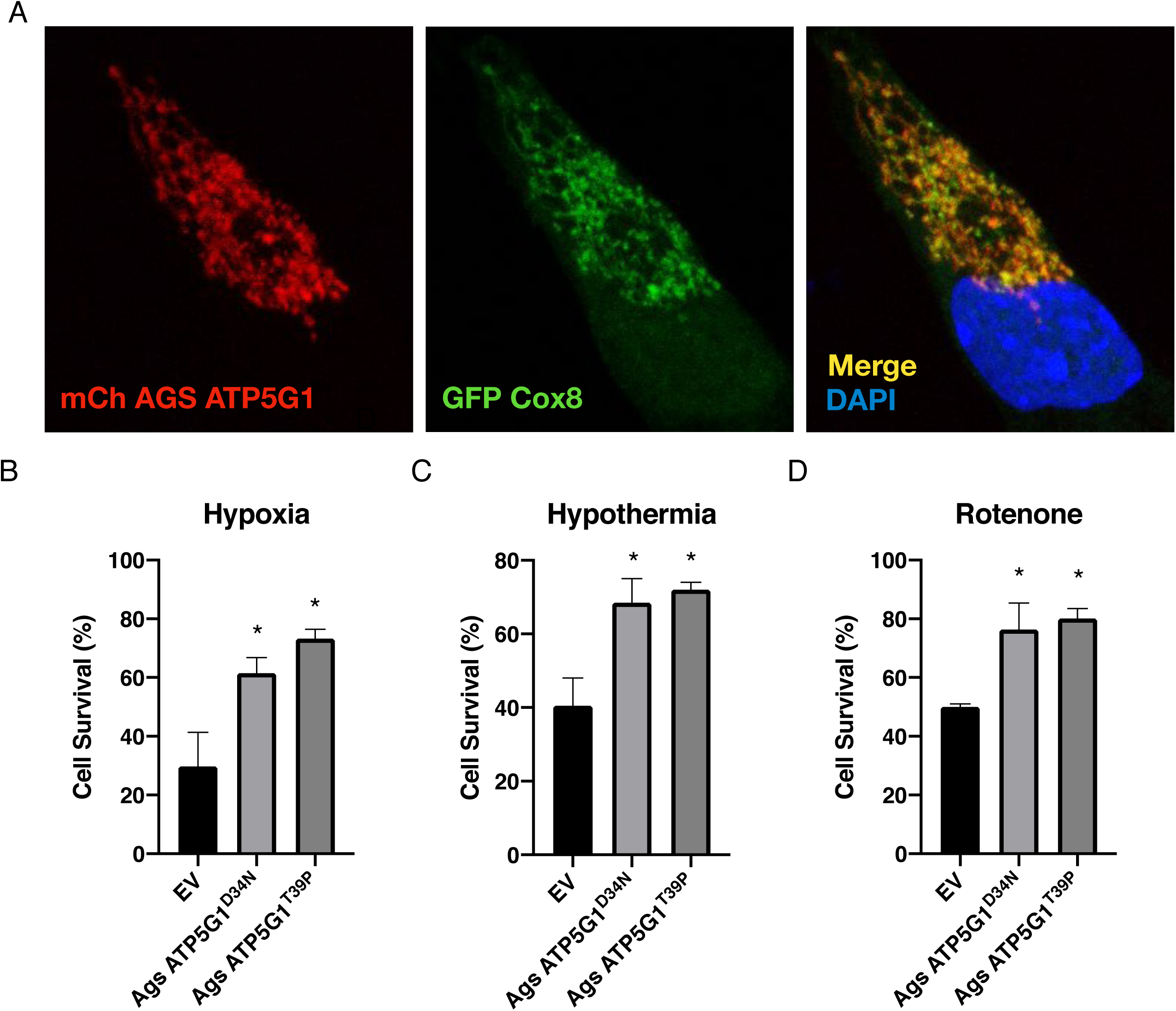
AGS ATP5G1 is targeted to mitochondria and overexpression of ATP5G1^D34N^ and ATP5G1^T39P^ mutant constructs do not alter cytoprotection. **(A.)** Confocal images of mouse NPCs expressing mCherry-AGS ATP5G1 and GFP-Cox8 demonstrating co-localization of the two proteins. **(B-D.)** Mouse NPCs expressing mutant AGS ATP5G1 isoforms (ATP5G1^D34N^, ATP5G1^T39P^) or empty vector (EV) and exposed to 24 hours of 1% O_2_ (B.), 31°C (C.), or 20mM rotenone (D.). Bar graphs represent data from 3 independent experiments with 3 replicates/genotype/condition, \**P*<0.05 vs EV.

**Figure 4 Supplement:**
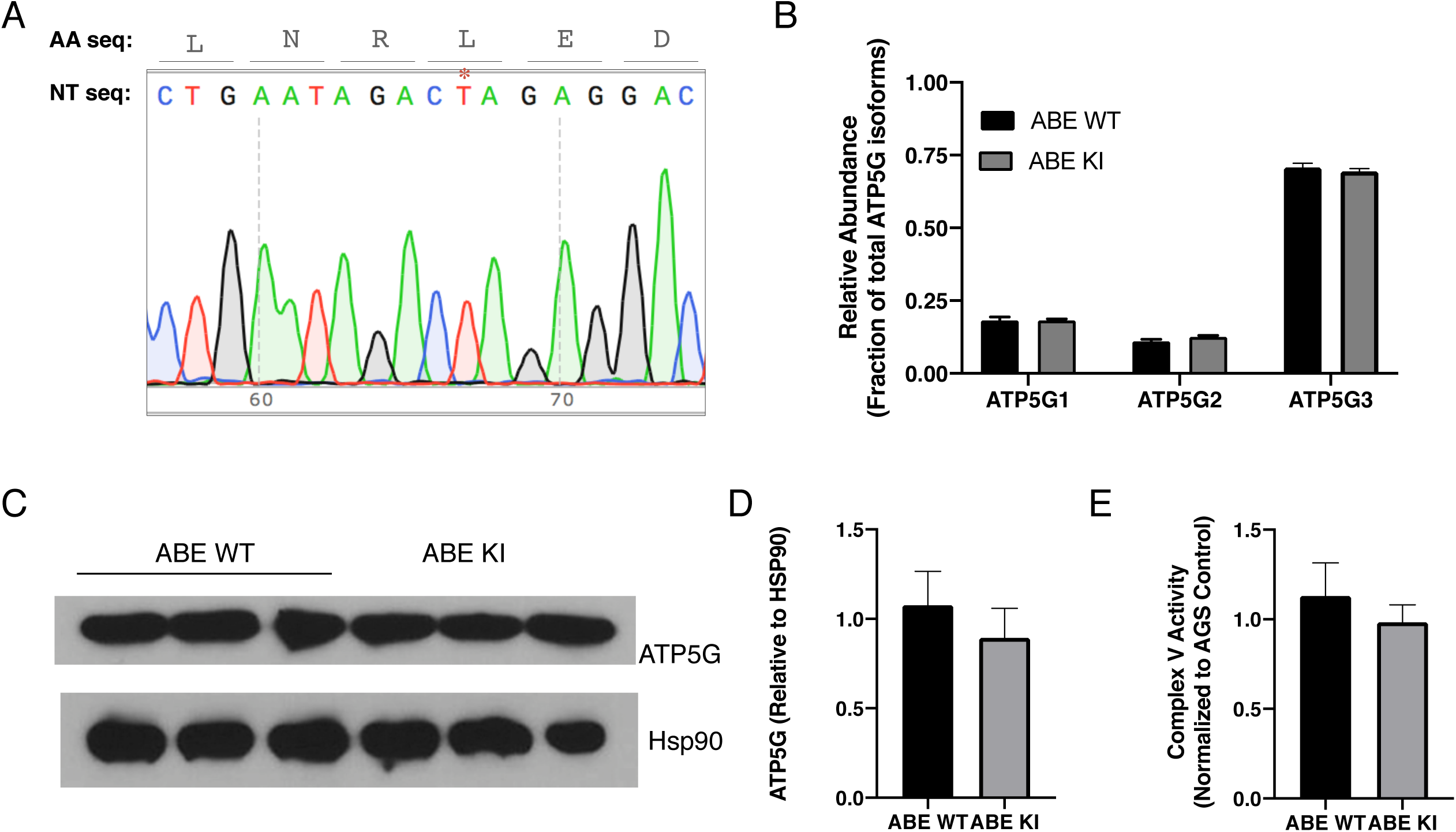
ATP5G expression in ABE cell lines. **(A.)** Sequencing data from an unsuccessfully edited clonal AGS cell line demonstrating the preservation of the wild-type sequence (* indicates wild-type thymidine). **(B.)** qRT-PCR for ATP5G1, ATP5G2, and ATP5G3 demonstrating the same relative abundance of ATP5G1 in ABE WT and ABE KI NPCs. Data from 3 independent experiments performed in triplicate. **(C.)** Representative western blot images demonstrating the relative abundance of ATP5G protein isoforms is similar in ABE WT and ABE KI cells. **(D.)** Quantification of western blots from 4 independent experiments with 2-3 replicates each. **(E.)** Complex V activity measured in mitochondrial extracts from ABE WT and ABE KI AGS NPCs. Data from 3 independent experiments with 3 replicates each.

**Supplemental Table 1.**
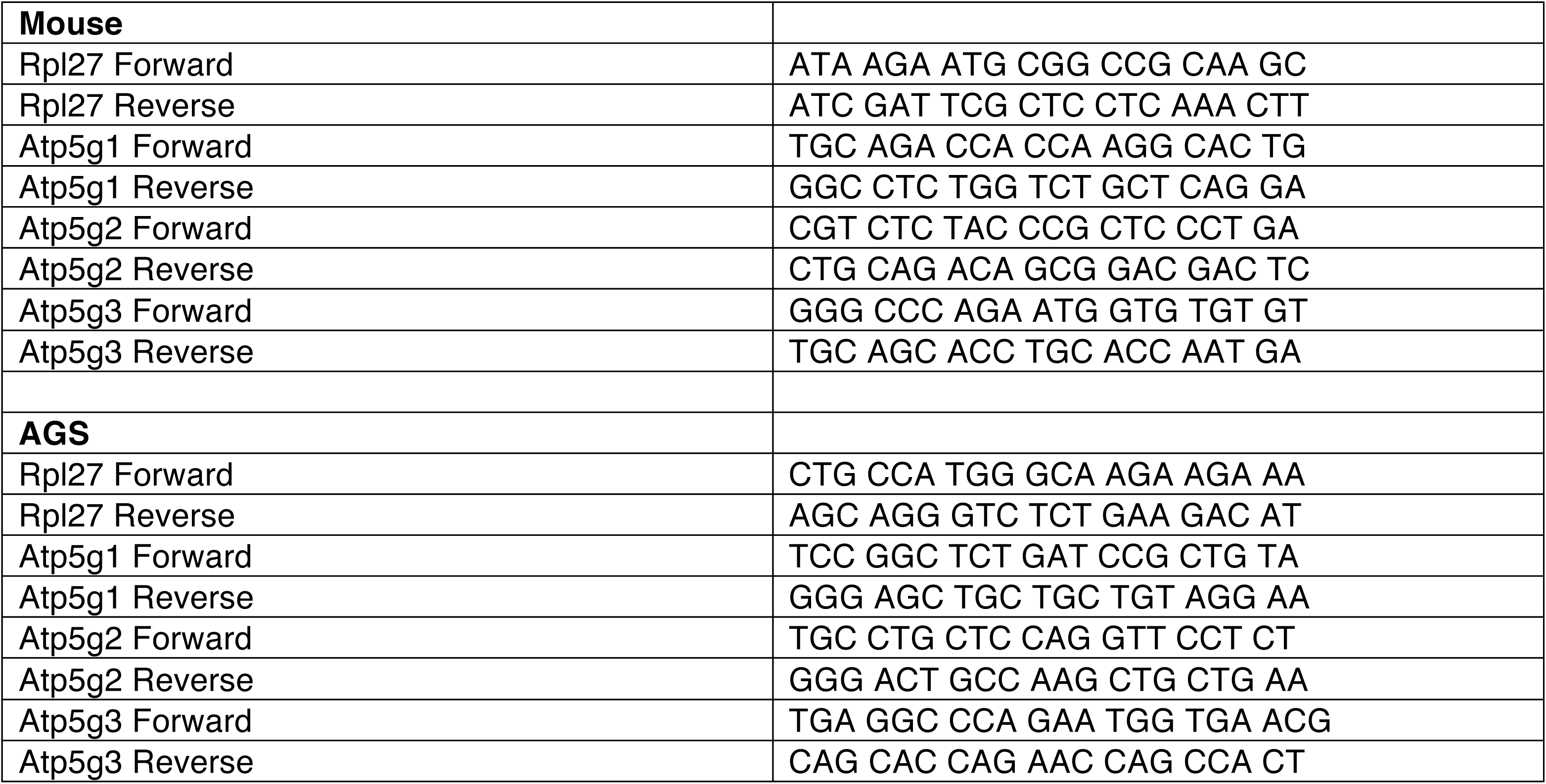
Species-specific primers used in quantitative RT-PCR.

## Notes

https://ucsf.box.com/s/wsetzswzvupx12sf5bjjxui5bnyn1hpu

https://ucsf.box.com/s/aelnc7c1hg2b7sen3cceatpm37nugllq

